# Pursuing reconsolidation in ethanol-CPP: memory reactivation in different conditions did not trigger destabilization

**DOI:** 10.1101/2024.08.31.610640

**Authors:** Flávia Zacouteguy Boos, Beatriz Déo Sorigotto, Cristiane Aparecida Favoretto, Fabio Cardoso Cruz, Isabel Marian Hartmann de Quadros

## Abstract

Consolidated memories can return to a labile state during retrieval, through destabilization, and must be reconsolidated to persist. Associative memories of contextual cues paired with hedonic effects of drugs of abuse exert a pivotal role in maintaining maladaptive behaviors in addiction. Thus, impairment of reconsolidation of drug-associated memories may provide a potential strategy to reduce drug-seeking and relapse in addiction. It is critical to understand the conditions under which a consolidated memory becomes labile and may undergo reconsolidation. After inducing ethanol Conditioned Place Preference (CPP; 2 g/kg ethanol, i.p.) in male mice, we examined different parameters during the reactivation session that could turn memory susceptible to disruption by systemic injection of the protein synthesis inhibitor cycloheximide (CHX, 100 mg/kg, i.p.). Reactivation with free access to the apparatus (similarly to a Test session, for 10, 5, or 3 min) or Reactivation sessions restricted to ethanol-paired compartment with no ethanol (for 10 or 5 min), or with the administration of a low dose of ethanol (5 min session), failed to reduced ethanol-preference after CHX administration. These findings suggest that boundary conditions constraint memory in ethanol-CPP to undergo reconsolidation.

## INTRODUCTION

The acquisition of associative memories plays a key role in predicting salient events, shaping behaviors in an adaptive way (Lee et al., 2017). However, different mental disorders involve formation of robust and persistent memories that guide undesirable or unhealthy behaviors, such as in post-traumatic stress disorders and drug addiction (Milton, 2013; Vaverková et al., 2020). Drug-associated memories, for example, have a pivotal role in triggering craving, drug-seeking, and relapse, which are major challenges for drug use disorders treatments (Milton & Everitt, 2012).

Conditioned place preference (CPP) is a well-known animal model to assess conditioned appetitive behaviors, rewarding effects of drugs and natural reinforcers (Bardo & Bevins, 2000; McKendrick et al., 2020). Increased preference (conditioned response) for a context (conditioned stimuli, CS) previously associated with the drug (unconditioned stimuli, US) indicates the formation of an associative memory between the drug rewarding effects and the drug-paired context (Tzschentke, 2007). Interventions targeting memory aim to disrupt or weaken these associations, thereby reducing the conditioned response (Milton, 2013).

Two main memory-related strategies to reduce conditioned response are extinction and reconsolidation (Rich & Torregrossa, 2018). Extinction learning is the most traditional procedure. It consists of a prolonged re-exposure to CS in absence of the US, until the CS no longer elicits the conditioned response (Vaverková et al., 2020). However, extinction has a limited therapeutic potential because extinction learning reflects a new memory trace that inhibits the previously one, resulting in behavioral inhibition rather than unlearning the association between context and drug effects. For instance, the original memory may return to guide behavior through reinstatement, spontaneous recovery, or renewal (Bouton, 2004), which can explain the relapse/reinstatement behavior after withdrawal.

By contrast, memory reconsolidation is a process that alters the original memory trace: it allows a consolidated memory to be destabilized through retrieval, requiring restabilization in order to persist (Nader et al., 2000; Lee et al., 2005). Because of that, interventions using reconsolidation process promote more efficient and enduring outcomes than extinction since they may prevent reinstatement in animal models. A retrieval session, also known as “reactivation”, is necessary to destabilize the original memory trace, leaving it susceptible to either strengthening (Inda et al., 2011; Forcato et al., 2014), weakening (Miller & Marshall, 2005; Nader et al., 2000), or updating (Pineyro et al., 2013; De Oliveira Alvares et al., 2013). In order to persist, memory should complete the restabilization phase. Different molecular mechanisms are required in restabilization, such as synthesis of novel proteins, and preventing such protein synthesis is a well-known strategy to disrupt reconsolidation (Nader et al., 2000; Milekic et al., 2006; Robinson & Franklin, 2007). The use of pharmacological (Nader et al., 2000; Taubenfeld et al., 2010; Valjent et al., 2006) or non-pharmacological (Goltseker et al., 2020; Haubrich et al., 2015; Popik et al., 2020) manipulations soon after or during reactivation session is used to modify (in its strength or content) or to disrupt memory. This approach has been explored also to modify drug-associated memories, in order to weaken the CS-US association and to reduce drug-seeking behaviors (Barak & Goltseker, 2021; Exton-McGuinness & Milton, 2018; Miller & Marshall, 2005; Milekic, 2006; Sorg, 2012; Xue et al., 2012).

However, reactivation session should be able to overcome boundary conditions that prevent memory from destabilization (Bustos et al., 2009; Pedreira et al., 2004; Robinson & Franklin, 2010; Exton-McGuinness & Milton, 2018). The occurrence of a prediction error, consisting of a mismatch between what was predicted by a consolidated memory and what actually happens during reactivation, is postulated as a key factor to bring memory again to this labile state (Exton-McGuinness et al., 2014; Fernández et al., 2016; Pedreira, 2004; Sevenster et al., 2013; Robinson et al., 2011). In this framework, different sources of novelty could promote error in prediction, such as the omission (Pedreira, 2004; Robinson et al., 2011) or delivery of different magnitudes of the US (Popik et al., 2020). The manipulation of these parameters during reactivation may facilitate memory destabilization.

Appropriate memory reactivation to trigger destabilization is more variable in CPP studies than in fear conditioning studies. Some data indicate a reduction in preference for different drugs of abuse when amnestic treatment was administered after reactivation with free access to the entire apparatus, similar to a test session (Miller & Marshall, 2005; Robinson & Franklin, 2010). However, in other labs this kind of reactivation did not induce reconsolidation, and adaptations in the protocol had to be done, such as re-exposure only to the drug-paired context (CS only; Lin et al., 2014) or an additional drug conditioning (CS-US; Valjent et al., 2006).

Ethanol is the most used drug worldwide and more than 100 million people follow the criteria for alcohol use disorder (WHO, 2018). In light of this, it is critical to investigate new therapies to prevent compulsive ethanol-seeking and relapse. Conditioned rewarding effects produced by ethanol are widely studied in the CPP paradigm (Cunningham & Prather, 1992; Groblewski et al., 2008; Macedo et al., 2018), although few of these studies investigate reconsolidation (Goltseker et al., 2020; Font & Cunningham, 2012; Lin et al., 2014).

In this paper, using CPP induced by ethanol, we manipulated different variables and parameters that could promote prediction error and possibly trigger a memory destabilization-reconsolidation process during a reactivation session.

## MATERIAL AND METHODS

### Subjects

Adult male swiss mice, 9-11 weeks old at the beginning of each experiment, were obtained from CEDEME (Centro de Desenvolvimento de Modelos Animais, Universidade Federal de São Paulo - UNIFESP). Mice were group housed (3-5 per cage) in acrylic cages (19 cm width x 30 cm length x 13 cm height), with free access to standard rodent chow and tap water. They were kept at an animal facility at the Departamento de Psicobiologia (CPP experiments) or the Departamento de Farmacologia (Inhibitory avoidance experiment, IA) (both at UNIFESP) with controlled temperature (21° ± 2°C) and humidity under 12 h: 12 h light / dark cycle (lights on at 7 h a.m.). The animals were acclimated for one week to the animal facility and they were handled for 4-5 days before experiments, for adaptation and stress reduction. All the procedures were approved by Ethics Committee on Animal Use (CEUA N° 9701271017) from UNIFESP.

### Drugs

All of the following drugs were injected intraperitoneally (i.p.):

Saline (NaCl 0.9%).

Ethanol (99.5% Synth®, Diadema, Brazil) was diluted in saline (15% w/v) and was injected at 13.3 ml/kg (2 g/kg dose) or 6,65 ml/kg (1 g/kg dose).

Cycloheximide (CHX, Sigma-Aldrich®, São Paulo, Brazil) was diluted in saline and was injected at 10 ml/kg (100 mg/kg dose; Murav’veva & Anokhin, 2007).

### Conditioned Place Preference (CPP)

#### Apparatus

The CPP experiments were conducted in two acrylic grey three-chamber apparatus (Insight Ltda, São Paulo, Brazil). Two distinct conditioning chambers (12.5 cm width x 15.5 cm length x 12 cm height) were connected by a small central one with gray walls and a smooth metal plate floor (12.5 cm width x 9.5 cm length x 12 cm height). One of the conditioning chambers had white walls with black horizontal stripes (2 cm thick) and a floor made of thin stainless-steel mesh. The other one had black walls with white vertical stripes (2 cm thick) and the floor was made of parallel stainless-steel bars (0,1 cm in diameter, spaced 0,4 cm apart). The time spent in the conditioning chambers, paired with saline or ethanol, was assessed using Ethovision XT® tracking software (Noldus, The Netherlands).

### CPP experimental procedure

Twelve animals per group were used at the beginning of experiments, based on previous studies of our research group. All animals were conditioned to ethanol 2 g/kg and saline at a corresponding volume. The CPP cage was cleaned with alcohol solution 70% in between sessions. An unbiased CPP procedure was used, and half of the animals received ethanol injections paired with the chamber with horizontal stripes, and the other half was paired with the chamber with vertical stripes. Treatment groups were semi-randomly assigned, so that different conditions were distributed within each home cage of mice (which CPP apparatus was used; which compartment was paired with ethanol; pharmacological treatment conditions), ensuring similar group averages in preference index during Preconditioning session.

General CPP procedure consisted of five phases: Preconditioning session (free exploration of entire CPP apparatus; 10 min), Conditioning sessions (restricted to the drug- or saline-paired compartment; 15 min), Reactivation session (which varied between the experiments), Test session (free exploration of entire CPP apparatus; 10 min), and Reinstatement session (free exploration of entire CPP apparatus; 10 min), as depicted in Figure 1A. During sessions in which animals could explore all the chambers in the CPP apparatus, each animal was placed in the central compartment, and then the guillotine doors were opened to allow access to ethanol- and saline-paired contexts. Conditioning sessions consisted of a total of eight sessions of 15 min distributed over four consecutive days, with two daily ethanol and saline conditioning sessions, separated by 6 h. The ethanol and saline conditioning sessions were alternated between morning and afternoon to avoid the association of ethanol with a specific time period (Ralph et al., 2013; Fig. 1A). Mice received the corresponding injection, and after 1-2 min were individually placed in ethanol or saline compartment, where they were confined for the duration of the session. Reactivation procedure took place 24 h after termination of conditioning sessions and the reactivation procedure differed between experiments, as detailed below. Immediately after memory Reactivation session, animals received an i.p. injection with saline or CHX and were returned to their home cage (except in Exp. 9). Conditioned preference was tested 24 h later. In experiments 1-3, 5-7, and 9, animals were given an ethanol priming injection (1 g/kg, i.p.) 5 minutes before the Reinstatement session to determine which memory trace was dominant at Reactivation session (Eisenberg et al., 2003). If extinction was the dominant memory trace, this would lead to preference reinstatement in the CHX group. Alternatively, if the dominant trace was the original memory and it was destabilized during the Reactivation session, the CHX group would maintain a low preference for the ethanol-paired compartment at Reinstatement session. Conditioned preference to ethanol was evaluated by a preference index, calculated by the time spent in the ethanol-paired context, divided by the sum of time spent in saline- and ethanol-compartments. All experiments were carried out during light cycle. Sessions with free exploration of entire CPP apparatus were carried out between 8 h and 13 h. Conditioning sessions took place between 8:30 h and 11:30 h, and between 14:30 h and 17:30 h.

**Figure 1.**
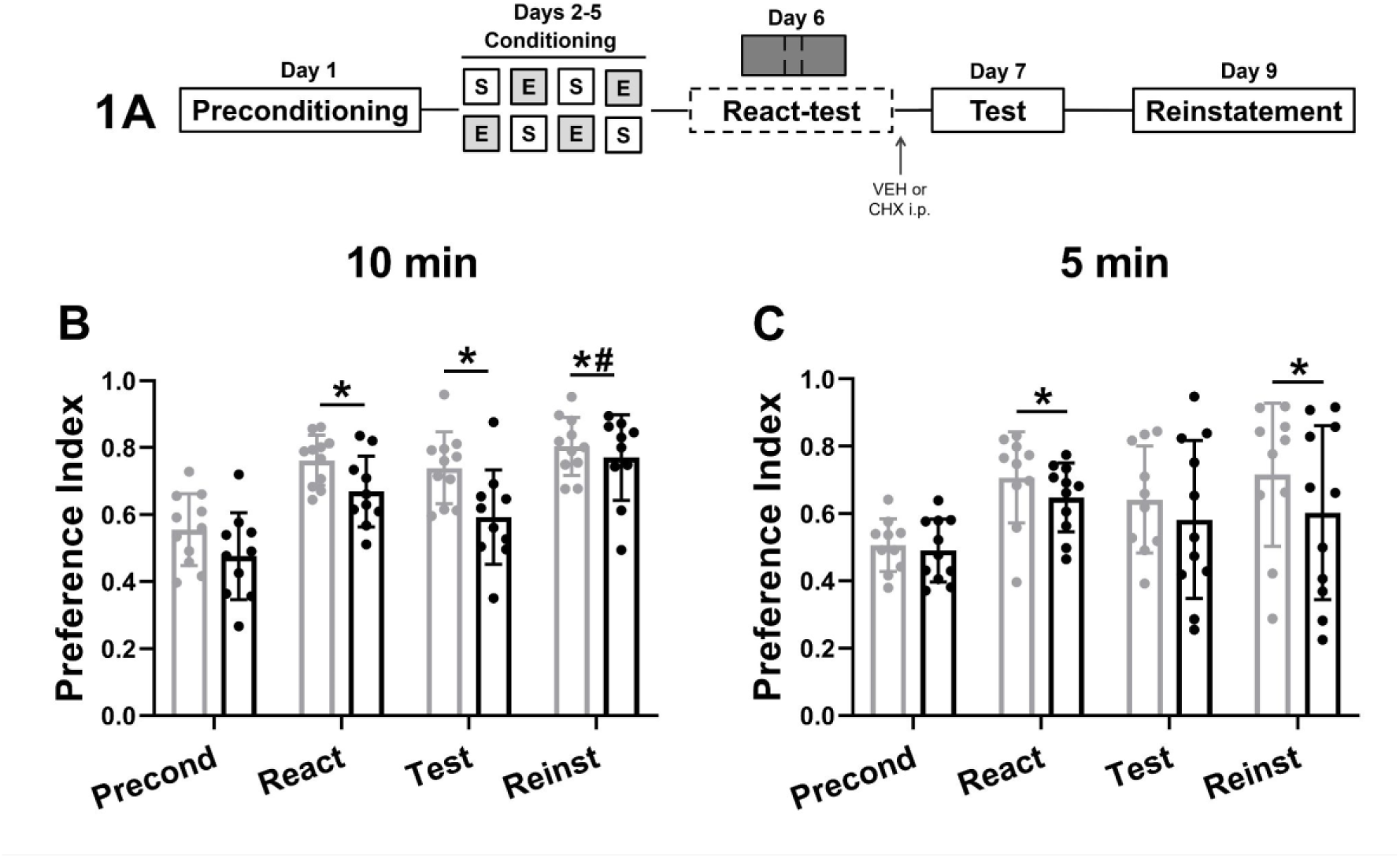
Preference index for ethanol-paired compartment during Preconditioning (Precond), Reactivation (React), Test and ethanol Reinstatement (Reinst) sessions, in mice that received CHX (100 mg/kg) or VEH (saline) i.p. right after the Reactivation session. (A) Experimental design. (B) Experiment 1: Ethanol-CPP using a 10 min Reactivation session with free access to the apparatus (React-test). (C) Experiment 2: Ethanol-CPP using a 5 min Reactivation session with free access to the apparatus (React-test). Data are represented by mean ± standard deviation. (*) p < 0.05 for session differences relative to Preconditioning. (#) p < 0.05 differences between Reinstatement and Test sessions.

### CPP Experimental Design

#### Reactivation with free access to the apparatus (similar to a test session, Reactivation-test)

A frequent condition used to effectively reactivate memory consists of recalling memory by exposing the animal to the entire apparatus, such as in a regular test (Bernardi et al., 2009; (Miller & Marshall, 2005; Robinson & Franklin, 2010). One day after the last Conditioning session, a Reactivation session was carried out in the same condition as the test (Reactivation-test), in which mice could freely explore all compartments for 10 min (Experiment 1), 5 min (Experiment 2), or 3 min (Experiments 3 and 4). Immediately after the Reactivation session, mice received vehicle or 100 mg/kg CHX i.p. and returned to their home cages. Twenty-four hours after Reactivation, ethanol-conditioned preference was assessed during a Test session, and 48 h later in a Reinstatement test, in which animals were challenged with a priming dose of ethanol 5 min before session (1 g/kg, i.p.) (except in Experiment 4). Conditioned preference was assessed in the Reactivation session and statistically analyzed in conjunction with Preconditioning, Test, and Reinstatement. The use of preference index, described above, allows to compare the conditioned preference between all sessions, even though the Reactivation session was shorter than the others (Exp. 2-4).

### Reactivation restricted to ethanol context (Reactivation-EtOH context)

In Experiments 5 and 6, a saline i.p. injection was administered to both groups 30 min before Reactivation session, as a control for future experiments. Reactivation restricted to ethanol context (Reactivation-EtOH context) consisted of re-exposure only to the ethanol compartment for 10 min (Exp. 5) or 5 min (Exp. 6). Immediately after, mice were injected with vehicle or 100 mg/kg CHX i.p. and returned to home cages. Preconditioning, Conditioning, Test, and Reinstatement sessions were identical as previously described.

### Reactivation restricted to ethanol context with a low dose of ethanol (Reactivation-EtOH context + EtOH)

In experiment 7, half of the dose of ethanol used during conditioning sessions (1 g/kg) was administered i.p. to mice 5 min before were confined to the ethanol-paired context for 5 min (Reactivation session). Soon after this Reactivation session, animals were injected with vehicle or 100 mg/kg CHX i.p. and returned to home cages. All other sessions of the experiment were the same as described before.

### Deconditioning-update approach: Two-contextual reactivations with a low dose of ethanol

Experiment 9 made use of a non-pharmacological strategy (with no CHX injection) to attenuate ethanol-conditioned preference in a deconditioning-update approach. This experiment was based on a study published by Popik et al. (2020) in auditory fear conditioning, in which they show that a weaker footshock during four reactivation sessions was able to reduce conditioned response in a subsequent test and this was mediated by reconsolidation. Then, the rationale of Experiment 9 is to update the motivational valence for ethanol-paired compartment by re-exposing each mouse to this compartment with administration of a lower dose of ethanol than that used in the Conditioning phase. We predicted that this procedure could promote a stronger negative prediction error, facilitating memory destabilization, and reducing the motivational valence (that Popik et al., 2020 called by deconditioning-update), resulting in reduced preference during the Test session. To test this hypothesis, Reactivation consisted of two sessions of re-exposure to ethanol context for 10 min, in two separate days (first in the morning and second in the afternoon), with an injection of a low dose of vehicle or ethanol (0,5 g/kg, i.p.) 5 min before each session. A group that did not undergo reactivation or receive any administration was included as a control. All other sessions were identical to previously described.

### Step-through Inhibitory Avoidance (IA)

#### Apparatus

The inhibitory avoidance was conducted in a room illuminated by red light. The apparatus consisted of an acrylic box divided into two compartments (21 cm width x 22 cm length x 21 cm height) separated by a horizontal sliding door. The floor of both compartments consisted of parallel stainless-steel bars (0.2 cm diameter, spaced 0.8 cm apart), connected by an electric shock generator (Ugo Basile Biological Research Apparatus, Italy). The “safe” compartment had white walls, whereas the “aversive” compartment had black walls.

### IA Experimental Procedure

Such as the other experiments, we initially planned to use 12 animals per group. However, with the high variability of responses during the Test session, we conducted a sample size calculation a *posteriori* to reach statistical power of 80%, and 14 animals per group were used. Groups were semi-randomly assigned in order to distribute different conditions within each home cage of mice (the number of shocks each animal received in Training session and the pharmacological treatment), ensuring similar Training strength across groups. The IA cage was cleaned with ethanol 70 % between sessions.

Mice were habituated to injection by administering saline i.p. (10 ml/kg) 48 h before Training. During all sessions, the animals were individually taken to the experimental room in a holding cage. After IA sessions, animals were housed temporarily in a different cage with cohabitants that were already trained or tested. It was used a multiple-trial procedure in Training session, which consisted in gently placing each mouse in the safe compartment for 90 s, when the sliding door was opened, giving access to the aversive compartment. When the mouse entered into the black, aversive compartment with its four paws, the door was closed 1 s later, and a 0.3 mA footshock was delivered for 1 s. This trial procedure was repeated until the mouse reached the learning criterion of avoiding crossing from the safe context to aversive compartment for 5 min (second trial) or 4 min (third trial). A maximum of three trials (three shocks) was used. The IA memory was tested 24 h later by placing each mouse in the safe compartment for 90 s, when the door was opened and the latency to cross to the dark compartment was registered for 9 min as a measure of memory retention. No footshock was delivered during the Test session. The latency was registered manually by the experimenter, who was blinded to the treatment condition. The experiment took place in the light cycle, between 9:00 h and 15:00 h. Experiment 8 was performed as a positive control for the amnestic effect of the 100 mg/kg CHX dose used in CPP experiments. To this end, animals were injected with vehicle or CHX 100 mg/kg i.p. immediately after the last trial in Training (Day 1). On the next day (Day 2), the memory was tested (Test) to assess the CHX effect on memory retention.

### Data analysis

In CPP experiments, animals which presented the preference index above or below the group mean ± 2 standard deviation (SD) at least in one free-exploration session were considered outliers and excluded from analysis (n = 17; 3 mice were excluded for health issues). In the CPP Experiments 1-4, we conducted two analyses. First, with all animals that were not outliers, and a second exploratory analysis was conducted only with those animals that reached a learning criterion, increasing at least 20% of preference index between Preconditioning and Reactivation sessions.

In both CPP and IA experiments, the data were analyzed by Generalized Estimating Equations (GEE), a statistical model designed to correlated data (repeated measures; Liang & Zeger, 1986). The main effects of session [CPP: Preconditioning, Reactivation (Exp. 1-4), Test, and Reinstatement; IA: Training and Test] and group (Exp. 1-8: vehicle and CHX; Exp. 9: No-Reactivation, Reactivation-Vehicle, and Reactivation-Ethanol), as well as the interaction between these factors, were evaluated for the conditioned ethanol-preference index (CPP), or latency to cross to the aversive compartment (IA). A covariation matrix with AR1 structure was used, and linear and gamma distribution were tested. Linear distribution was the best fit to the data for all CPP experiments, whereas gamma distribution was the best one for IA experiment, based on the lowest value of Quasi Likelihood Under Independence Model Criterion (QIC). When GEE detected significant effects, Sidak *post-hoc* was used to find pairwise comparisons. Generalized Linear Models (GzLM) were used to compare the number of shocks between groups in IA experiment; linear distribution was the best fit based on the lowest Akaike’s Information Criterion (AIC). All analyses were carried out using IBM SPSS Statistics for Windows, version 20 (IBM Corp., Armonk, N.Y., USA), and all graphs were made by GraphPrism 9 (GraphPad Software Inc., La Jolla, CA, USA). A p value ≤ 0.05 was adopted as a criterion to consider statistically significant differences. The data are shown as mean ± standard deviation (SD; CPP experiments) or median ± 95 % of confidence interval (IA experiment).

## RESULTS

### 1. Reactivation with free access to the apparatus (similar to a test session, Reactivation-test)

In this series of experiments (1-4), after the ethanol/saline conditioning sessions, mice were exposed to a Reactivation session using a “testing condition”, when they were allowed to freely explore the CPP apparatus for different periods of time, as specified below. Right after the Reactivation session, mice received CHX or saline (VEH) i.p. and were tested for ethanol induced CPP 24h later. A Reinstatement session was carried out after the injection of a lower dose of ethanol (except in Exp. 4, due to molecular brain analyses that had been planned).

#### 1.1. Reactivation-test - 10 min session

In Experiment 1, GEE detected session (Wald = 97.201, DF = 3, p < 0.001) and group effect (Wald = 10.963, DF = 1, p = 0.001), but not session by group interaction (Wald = 3.064, DF = 3, p = 0.382; n VEH = 11, n CHX = 10; Fig. 1B). Animals acquired and maintained preference for the ethanol-paired context across all sessions, when compared to Preconditioning (p < 0.001). There was no decrease in preference during the Test session, relative to the Reactivation session (p = 0.226), and a lower dose of ethanol further increased conditioned preference for ethanol-paired context, during a Reinstatement test (Test vs. Reinstatement, p = 0.002).

In order to reduce variability in ethanol induced CPP, we conducted a separate analysis considering only animals that increased at least 20% of preference during the Reactivation session, relative to Preconditioning test (n = 8 per group). GEE detected the same pattern of differences, with session (Wald = 325.155, DF = 3, p < 0.001) and group effect (Wald = 6.800, DF = 1, p = 0.009), but no interaction between factors (Wald = 2.570, DF = 3, p = 0.463; data not shown).

These results indicate that a 10 min session of Reactivation with free access to the apparatus was not able to trigger destabilization-reconsolidation in ethanol-CPP.

#### 1.2. Reactivation-test - 5 min session

In experiment 2, we tested if a 5 min Reactivation session with free access to the apparatus was able to destabilize the ethanol-conditioning memory (n VEH = 10, n CHX = 11). GEE analysis indicated only session effect (Wald = 54.839, DF = 3, p < 0.001) with no significant effects of group or group X session interaction (group: Wald = 1.816, DF = 1, p = 0.178; interaction: Wald = 1.629, DF = 3, p = 0.653). There was a conditioning effect, shown by an enhanced preference for ethanol-paired context during Reactivation session (p < 0.001 relative to Preconditioning). During the Test, the ethanol-conditioned preference did not persist; however, this preference was recovered during the Reinstatement test following an ethanol challenge, as shown by the comparison to Preconditioning (p = 0.070 for Test and p = 0.015 for Reinstatement; Fig. 1C).

Analysis including only animals that increased at least 20% of preference in Reactivation relative to the Preconditioning session, only detected a significant session effect (data not shown; Wald = 151.639, DF = 3, p < 0.001; group: Wald = 1.757, DF = 1, p = 0.185; interaction: Wald = 1.070, DF = 3, p = 0.784; n = 8 per group).

These results show that a 5 min session of Reactivation using test conditions failed to induce destabilization in ethanol-CPP.

#### 1.3. Reactivation-test - 3 min session

In Experiment 3 we tested an even shorter Reactivation session, lasting 3 min in the same reactivation condition (n VEH = 12, n CHX = 11). GEE analysis detected only a significant session effect (Wald = 63.623, DF = 3, p < 0.001; group: Wald = 0.187, DF = 1, p = 0.665; interaction: Wald = 4.144, DF = 3, p = 0.246). There was acquisition and maintenance of conditioned preference for the ethanol context, indicated by differences between Preconditioning and all other sessions (p ≤ 0.005; Fig. 2B). There was a reduction in ethanol-preference index between Reactivation and Test sessions (p = 0.003), and Reinstatement session did not differ from Reactivation or Test sessions (p > 0.05).

**Figure 2.**
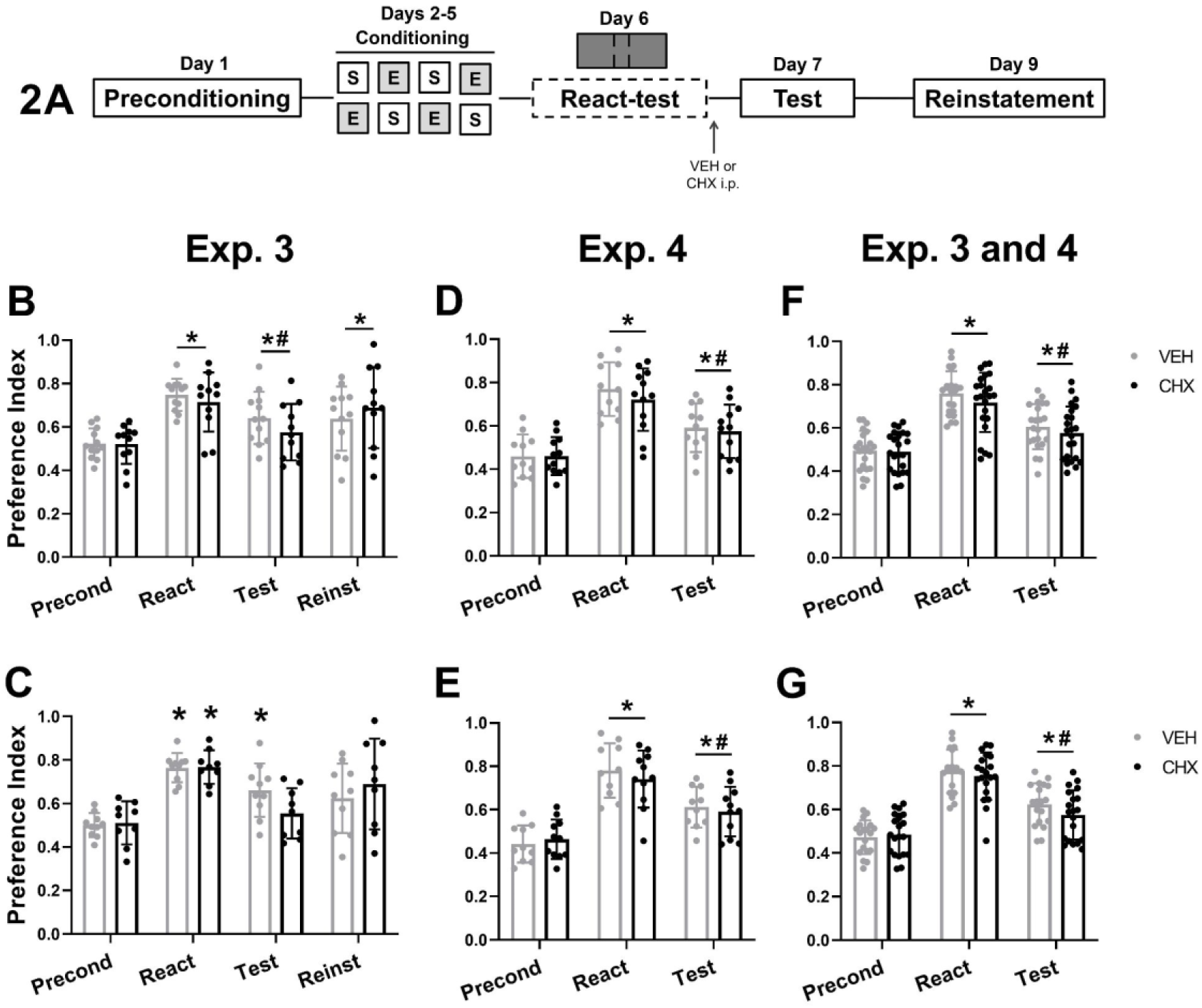
Preference index for ethanol-paired compartment during Preconditioning (Precond), Reactivation (React), Test, and ethanol Reinstatement (Reinst) sessions, in mice that received CHX (100 mg/kg) or VEH (saline) i.p right after a 3 min Reactivation under test condition session. (A) Experimental design. (B) Experiment 3: ethanol-CPP using a 3 min Reactivation with free access to the apparatus (React-test). (D) Experiment 4: replication of Experiment 3 with a new batch of animals. (F) Pooled the data from Experiment 3 and 4. (C), (E) and (G): Subsets of animals from Experiments 3 (C), 4 (E), and from both experiments (G), which had increased at least 20 % of ethanol-conditioned preference during Reactivation session, relative to Preconditioning. Data are represented by mean ± standard deviation. (*) p < 0.05 for session differences relative to Preconditioning. (#) p < 0.05 differences between Reinstatement and Test sessions.

The GEE analysis with only those animals that increased at least 20 % of preference for the ethanol context during Reactivation session relative to Preconditioning, detected a significant effect of session (Wald = 298.496, DF = 3, p < 0.001) and interaction between session and group (Wald = 13.375, DF = 3, p = 0.004; group: Wald = 0.042, DF = 1, p = 0.837; n VEH = 10, n CHX = 9). *Post-hoc* Sidak for interaction indicated that the groups acquired conditioned preference for ethanol (Reactivation relative to Preconditioning: p < 0.001). Treatment with CHX reduced ethanol-preference during the Test (compared to Reactivation session, p < 0.001), although there was no difference between groups in this session (p = 0.680; Fig. 3C). A lower dose of ethanol failed to reinstate ethanol-preference in the CHX group during Reinstatement (relative to Preconditioning: p = 0.398). The control group (VEH) showed significant preference for the ethanol-context during Reactivation and Test sessions (p ≤ 0.005), but no significant preference during Reinstatement test (p = 0.333). The decreased preference in CHX group at the Test session corroborates the hypothesis that 3 min Reactivation-test induced memory sensitive to disruption, but the lack of difference between groups in this session leaves doubts about the effectiveness of this protocol.

**Figure 3.**
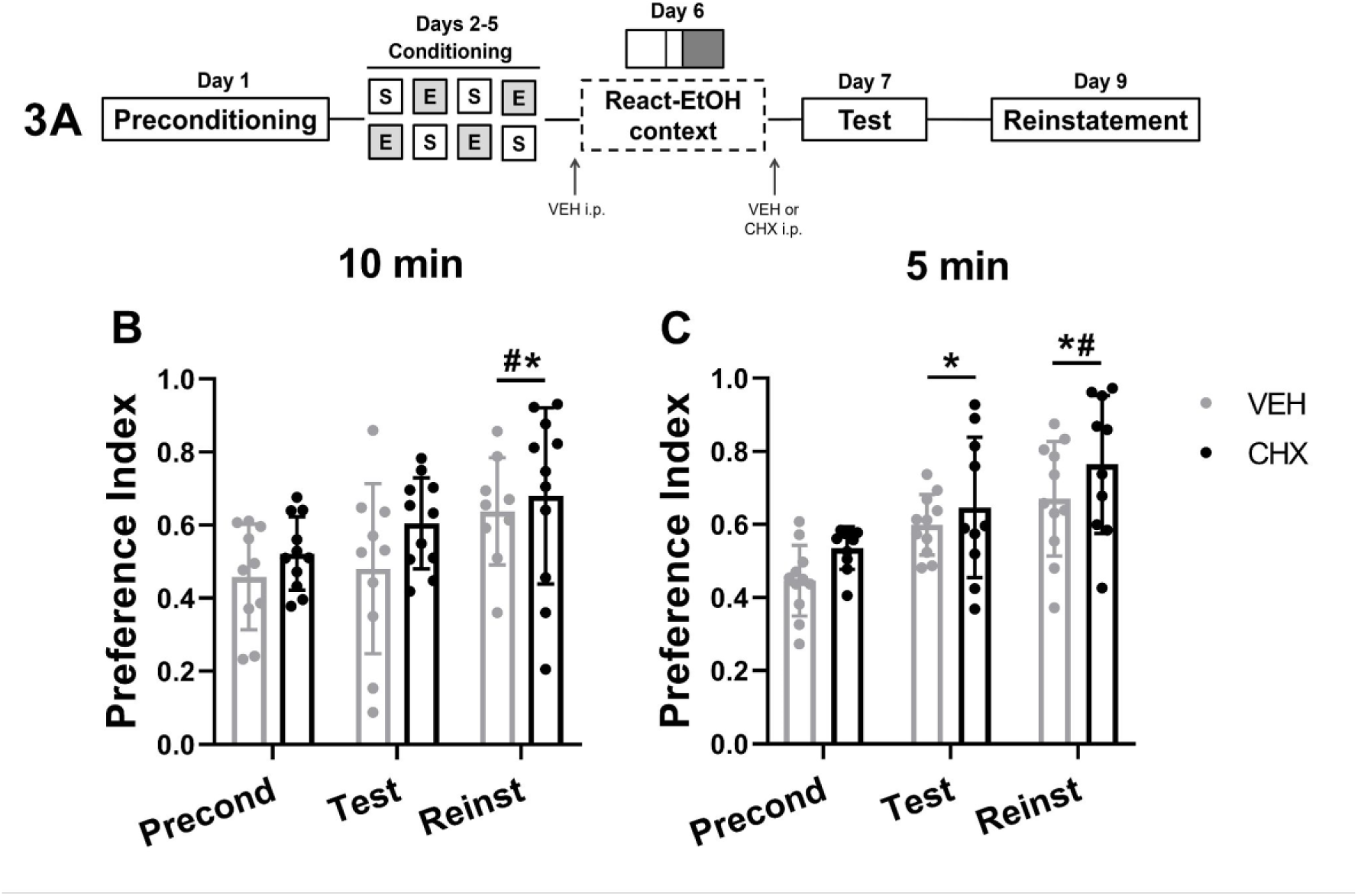
Preference index for ethanol-paired compartment during Preconditioning (Precond), Test and ethanol Reinstatement (Reinst) sessions, in mice that received CHX (100 mg/kg) or VEH (saline) i.p. right after the Reactivation session. (A) Experimental design. (B) Experiment 5: Ethanol-CPP using a 10 min Reactivation session restricted to the ethanol context condition (React-EtOH context). (C) Experiment 6: Ethanol-CPP using a 5 min Reactivation session restricted to ethanol-paired context condition. Data are represented by mean ± standard deviation. (*) p < 0.05 for session differences relative to Preconditioning. (#) p < 0.05 differences between Reinstatement and Test.

To clarify this issue, the experimental design was replicated in a new batch of animals (Experiment 4), in order to collect brain tissue for molecular analysis after the Test, if it was confirmed that CHX impaired reconsolidation. However, GEE analysis detected only session effect (Wald = 92.200, DF = 2, p < 0.001), and no effect of group (Wald = 0.464, DF = 1, p = 0.496) neither session by group interaction (Wald = 0, 719, DF = 2, p = 0.698; n VEH = 11, CHX = 12). *Post-hoc* Sidak for the session effect indicated that animals increased preference during Reactivation (p < 0.001), with a reduction during Test (React vs. Test: p < 0.001), although there was still difference between Test and Preconditioning (p < 0.001; Fig. 2D). Analyzing the data of mice with a minimal 20% increase in preference for the ethanol-context during Reactivation relative to Preconditioning, the results remained similar to previous analysis, with only a significant effect of session (Wald = 117.616, DF = 2, p < 0.001; group: Wald = 0.167, DF = 1, p = 0.683; interaction: Wald = 1.270, DF = 2, p = 0.530; n VEH = 10, n CHX = 11) and results are depicted in Fig. 2E.

Pooling together the data from Experiments 3 and 4, there was only a significant session effect (Wald = 131.180, DF = 2, p < 0.001; group: Wald = 1.569, DF = 1, p = 0.210; interaction: Wald = 0.826, DF = 2, p = 0.662; n VEH = 22, n CHX = 23), and the *post-hoc* detected a significant difference between all sessions (p < 0.001; Fig. 2F). The analysis including only mice that increased at least 20% preference on Reactivation relative to Preconditioning, confirmed, again, only an effect of session (Wald = 253.082, DF = 2, p < 0.001; group: Wald = 0.821, DF = 1, p = 0.365; interaction: Wald = 3.067, DF = 2, p = 0.216). All sessions differed from one another (p < 0.001; Fig. 2G; n VEH = 19, n CHX = 20).

Thus, despite an apparent promising outcome in experiment 3, together these data suggest that using a reactivation session of 3 min with free access to the apparatus was not able to destabilize ethanol-CPP memory.

### Reactivation restricted to ethanol context (Reactivation-EtOH context)

In this section, destabilization of ethanol-conditioned preference was attempted by using reactivation sessions (10 min or 5 min) in which mice were restricted to the ethanol context, in the absence of the drug (reward omission; Lin et al., 2014). Immediately after the Reactivation session, mice received VEH or CHX i.p..

#### 2.1. Reactivation-EtOH context - 10 min session

GEE analysis detected only session effects in Experiment 5 (Wald = 20.043, DF = 2, p < 0.001; group: Wald = 1.969, DF = 1, p = 0.161; interaction: Wald = 0.988, DF = 2, p = 0.610; n VEH= 10, n CHX = 11). *Post-hoc* Sidak revealed that ethanol-conditioned preference was only expressed during Reinstatement (compared with Preconditioning, p < 0.001; compared with Test, p = 0.005; Fig. 3B) but not during the Test session. In this experimental design, ethanol-related memory was not assessed before or during reactivation session.

Thus, a 10 min reactivation session in the ethanol-paired compartment failed to induce the expected destabilization of ethanol-conditioned preference, given the lack of CHX treatment effect.

#### 2.2. Reactivation-EtOH context - 5 min session

GEE analysis of Experiment 6 also indicated only a session effect (Wald = 34.780, DF = 2, p < 0.001; group: Wald = 3.286, DF = 1, p = 0.070; interaction: Wald = 0.725, DF = 2, p = 0.696) and the results are illustrated in Figure 3C (n VEH = 11, n CHX = 10). *Post-hoc* Sidak detected a significant difference among all sessions, indicating that there was an increase in preference for the ethanol context during Test (p < 0.001) and a further increment in preference during the Reinstatement session (Test vs. Reinstatement: p = 0.012), demonstrating that ethanol challenge further increased ethanol preference.

These results together suggest that a 10 min or a 5 min Reactivation session, restricted to the ethanol context but with omission of drug, were not able to induce reconsolidation.

### 3. Reactivation-EtOH context + EtOH – 5 min

In this experiment, a 5 min reactivation session was also restricted to the ethanol context. A lower dose of ethanol (1 g/kg i.p.) was administered to mice 5 min before Reactivation session, attempting to achieve memory destabilization by adding a dose-related prediction error. The GEE detected an effect of session (Wald = 92.045, DF = 2, p < 0.001), but not of group (Wald = 0.060, DF = 1, p = 0.806) or interaction between these factors (Wald = 0.221, DF = 2, p = 0.895; n VEH = 12, n CHX = 11). *Post-hoc* Sidak detected a significant increase in ethanol-preference during Test and Reinstatement sessions, relative to Preconditioning (p < 0.001; Fig. 4B).

**Figure 4.**
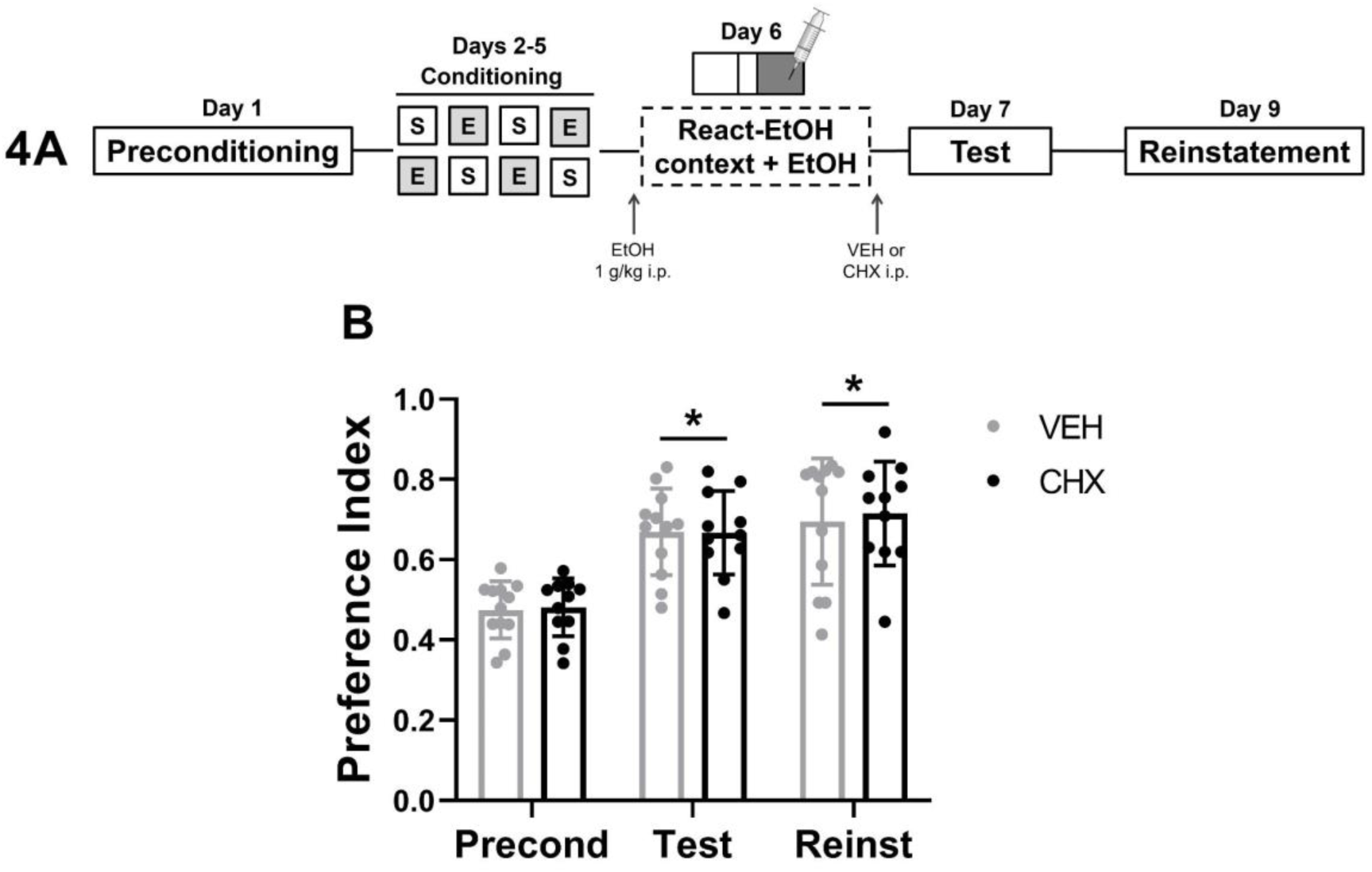
Preference index for ethanol-paired context during Preconditioning (Precond), Test and ethanol Reinstatement (Reinst) sessions, in mice that received CHX (100 mg/kg; i.p.) or VEH (saline i.p.) right after the Reactivation restricted to ethanol-paired context for 5 min, under a low dose of ethanol (1 g/kg; React-EtOH context + EtOH). (A) Experimental design. (B) Experiment 7: Preference for ethanol-context during Preconditioning (Precond), Test, and Reinstatement (Reinst) sessions. Data are represented by mean ± standard deviation. (*) p < 0.001 for session differences compared with Preconditioning.

This result indicates that a reactivation session restricted to the ethanol-context with half the dose of ethanol used during Conditioning sessions, failed to induce reconsolidation.

### 4. CHX 100 mg/kg post-training disrupted memory consolidation in Inhibitory Avoidance task

In this experiment, we wanted to ensure that the CHX dose used in CPP experiments, was an effective memory-disrupting dose. For that, we used a post-training administration of CHX (100 mg/kg, i.p. or vehicle, n = 14/group) in the Inhibitory Avoidance task (IA), as a positive control. Different from CPP paradigm, IA requires only a single day of learning and that is why this paradigm was chosen.

GEE analysis detected the effect of session (Wald = 46.733, DF = 1, p < 0.001), group (Wald = 8.256, DF = 1, p = 0.004), and session by group interaction (Wald = 8.599, DF = 1, p = 0.003). Both groups increase latency to cross to the aversive context during the Test compared to Training session (VEH p < 0.001, CHX p = 0.018; Fig. 5B). However, during the Test, mice treated with CHX presented a reduced latency to cross to the aversive context, relative to the VEH group (p = 0.021). This result cannot be attributed to the intensity of Training because there was no difference between groups relating to the number of shocks received in this session (Wald = 0.083, DF = 1, p = 0.773; Fig. 5C). This result support that 100 mg/kg of CHX is a suitable dose to induce amnestic effects.

**Figure 5.**
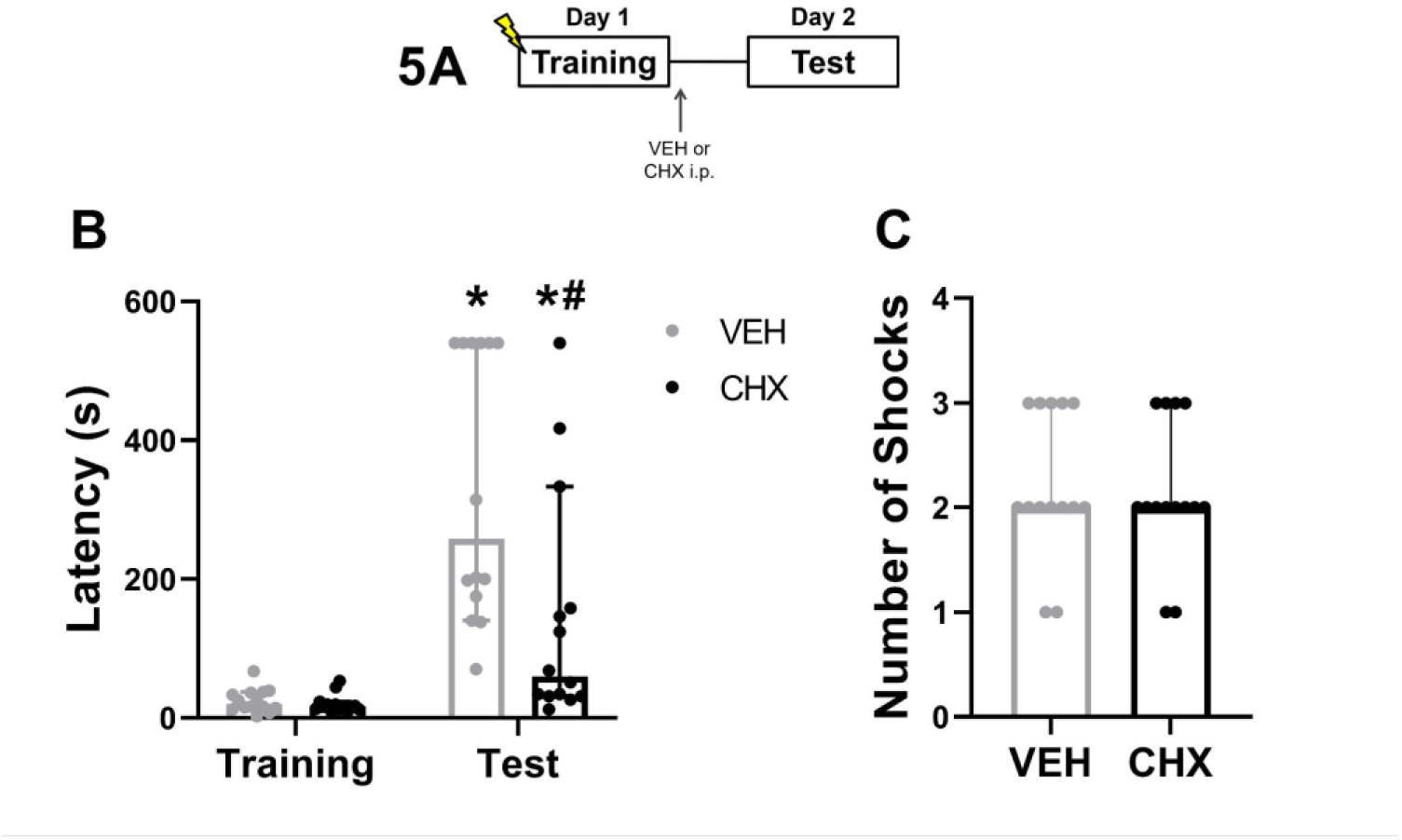
Time latency (s) to cross from safe to aversive compartment during Training and Test sessions in the Inhibitory Avoidance task, in mice which received CHX (100 mg/kg; i.p.) or VEH (saline i.p.) right after the Training session. (A) Experimental design. (B) Latency to enter in the shock context in Training and Test sessions. (C) Number of shocks during Training for each group. Data are represented by median ± confidence interval. (*) p < 0.05 for differences between Training and Test for each group. (#) p = 0.021 for difference between groups in Test session.

### 5. Deconditioning-update approach: Two contextual reactivations with a low dose of ethanol

In Experiment 9, we investigated whether two 10 min re-exposure sessions to the ethanol compartment, under the effect of an even low dose of ethanol (0.5 g/kg), would be able to update memory by reducing its motivational valence, and diminishing preference for this context in the Test session. To this end, a day after the Conditioning phase mice were divided into three groups: one containing the animals that stayed in home cage without any drug administration (No Reactivation group, n = 10), and two others that had their memory reactivated, one of them receiving saline (React – VEH, n = 11) and the other one ethanol (React – EtOH, n = 11) i.p. 5 min before the Reactivation sessions. GEE analysis detected an effect of session (Wald = 63.924, DF = 2, p < 0.001) and group (Wald = 7.374, DF = 2, p = 0.025), but not session by group interaction (Wald = 5.545, DF = 4, p = 0.236). The *post-hoc* Sidak for session detected a difference between all sessions (p < 0.001), with increased preference during Test and ethanol challenge further increasing ethanol-conditioned preference retrieval on Reinstatement (Fig. 6B). The *post-hoc* Sidak for group indicated that the No Reactivation group showed a greater preference for the ethanol context than the React – VEH group (p = 0.020).

**Figure 6.**
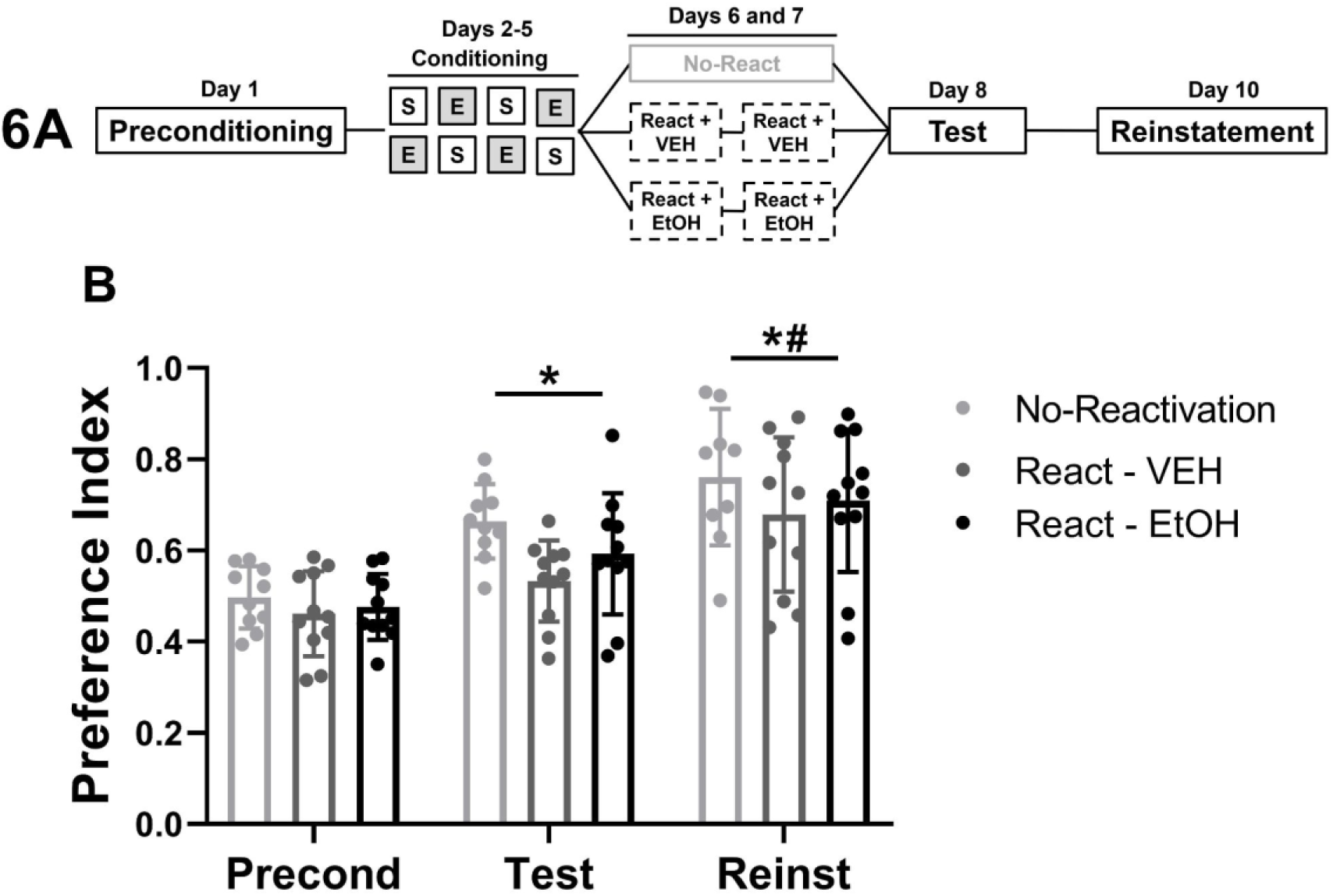
Preference index for ethanol-paired context over sessions in mice which had undergone two 10 min Reactivations restricted to ethanol-paired compartment under the effect of a very low dose of ethanol (EtOH 0.5 g/kg, i.p.) or saline (VEH; Deconditioning-update, Exp. 9). (A) Experimental design. (B) Experiment 9: conditioned preference for ethanol compartment during Preconditioning (Precond), Test, and Reinstatement (Reinst) sessions. (*) p < 0.05 for session differences relative to Preconditioning. (#) p < 0.05 for differences between Reinstatement and Test.

Therefore, two-contextual reactivations with a low dose of ethanol were unable to reconsolidate ethanol-CPP memory, with no indication of memory deconditioning.

## DISCUSSION

In this study, we addressed whether different levels of prediction error during retrieval are effective to trigger destabilization in memory underlying ethanol conditioned place preference. We specifically manipulated the reactivation session (condition and duration) that could have rendered memory destabilization and allowed for CHX treatment to effectively disrupt reconsolidation in the ethanol-CPP. Additionally, we tested a non-pharmacological manipulation to investigate whether this memory could be updated through reconsolidation, reducing its motivational valence (deconditioning-approach), by using a lower magnitude of US in combination with re-exposure to CS-paired with ethanol. However, despite the different approaches and parameters tested in this series of experiments, we were not able to overcome the boundary conditions of memory underlying conditioned preference, and thus ethanol preference memory was not changed.

Memory reconsolidation is a conserved evolutionary phenomenon (Lee et al., 2017) that has been largely investigated in a range of behavioral paradigms, memory types (Exton-McGuinness & Milton, 2018; Milekic et al., 2006; Nader et al., 2000; Myskiw et al., 2008), and species including humans (Milekic & Alberini, 2002); Pedreira, 2004; Sevenster et al., 2013). A postulated role for reconsolidation is to update the memory and to maintain its predictive role in a constantly changing environment (Lee et al., 2017). In order to be reconsolidated, memory must be first destabilized, and it is usually triggered by cue reminders which allow memory reactivation through retrieval. Retrieval is necessary to induce reconsolidation, but it is not sufficient on its own (Bustos et al., 2009; Robinson & Franklin, 2010; Suzuki et al., 2004; Valjent et al., 2006; but see Delorenzi et al., 2014). Boundary conditions prevent memory to return to a labile state, when it would be susceptible to modifications (Robinson & Franklin, 2010; Suzuki et al., 2004). These conditions encompass features of memory itself, such as its age (Milekic & Alberini, 2002; Robinson & Franklin, 2010) and its strength (Bustos et al., 2009; Liddie & Itzhak, 2016; Suzuki et al., 2004), and conditions during memory retrieval, which could be manipulated to overcome the boundary conditions imposed by the nature of memory (Exton-McGuinness & Milton, 2018).

Studies with conditioned fear memories paradigms have contributed largely for reconsolidation literature. However, not all psychological features and neurobiological mechanisms of these memories are shared with appetitive ones (Itzhak et al., 2014; Young et al., 2016). Compared to fear conditioning, appetitive conditioning memories require additional CS-US pairings to be efficiently stored (Itzhak et al., 2014). Increasing the magnitude of US (e.g., intensity of shock or drug dose) and/or the number of CS-US pairings tend to elevate expression of the conditioned response (Risinger & Oakes, 1996; Cunningham & Prather, 1992) until an asymptotic and stronger learning is reached.

Given that in our study we conditioned mice to ethanol using four conditioning drug sessions using an intermediate to high dose (2 g/kg i.p.; Groblewski et al., 2008), it is possible that a robust ethanol-context association was acquired, contributing to a conditioned preference memory that is resistant to reconsolidation (Bustos et al., 2009; Liddie et al., 2016; Robinson & Franklin, 2010; Suzuki et al., 2004). This notion agrees with CPP studies in which stronger conditionings (using escalating doses of cocaine across conditionings: Liddie et al., 2016; or more CS-morphine pairings: Robinson & Franklin, 2010) requires specific strategies to conditioned preference memory undergoes reconsolidation. However, in our study different reactivation session conditions reverted, or partially reduced, the conditioned preference at the test session (Exp. 2-5). It is unlikely that the resistance to destabilization was due to the strength of ethanol-CPP memory. In addition, memory age does not seem to interfere with reconsolidation in our protocol, as conditioned memory was reactivated 24 h after the last conditioning session.

Specific strategies are required by each memory paradigm to overcome boundary conditions. Despite these specificities, many authors posit that a mismatch between expected outcomes and current events during the retrieval session is necessary. This mismatch is known as a prediction error (Exton-McGuinness et al., 2014; Fernández et al., 2016; Sevenster et al., 2013). Furthermore, contextual reactivation seems to be necessary to trigger destabilization, since amnestic treatments without re-exposure to drug-paired context or to the whole of CPP box did not engage reconsolidation (Bernardi et al., 2009; Brown et al., 2008; Lin et al., 2014; Taubenfeld et al., 2010; Théberge et al., 2010; Valjent et al., 2006; (Yu et al., 2013).

In our study, all reactivation sessions were contextual and should be able to induce some degree of prediction error theoretically. Possibly, reactivation with free access to the apparatus is the less likely to promote high levels of prediction error, because mice have already explored the whole apparatus in a drug-free state during Preconditioning. Nevertheless, many studies using this condition were successful in disrupting memory reconsolidation in CPP (cocaine-CPP: Bernardi et al., 2009; Miller & Marshall, 2005; Théberge et al., 2010; morphine-CPP: Robinson et al., 2011), and in preventing the return of conditioned preference by drug priming injection (morphine-CPP: Yu et al., 2013). However, this type of reactivation does not constitute a universal condition to trigger reconsolidation, since other studies failed to show effect of amnestic treatments after this type of reactivation for 15 min (ethanol-CPP: Font & Cunningham, 2012; morphine-CPP: Yim et al., 2006) or 10 min (morphine-CPP: Milekic et al., 2006). These last reports agree with ours, as our data showed that neither 10 min, 5 min, nor 3 min of reactivation with free access to the apparatus (React-test) were able to make ethanol-CPP memory susceptible to CHX disruption.

In our study, confinement to the drug context was experienced only accompanied by an ethanol injection during conditioning sessions. Thus, we predicted that a reactivation session restricted to drug-paired context with no ethanol administration (CS only), or with a low dose of ethanol (CS-weakerUS), could increase the prediction error by violating the expectancy to receive the US, which is pointed as an important factor to induce destabilization (Pedreira et al., 2004; Fernández et al., 2016). Fernández et al., 2016). In contrast to other studies with morphine-, cocaine-, and ethanol-CPP (De Carvalho et al., 2014; Escosteguy-Neto et al., 2016; Lin et al., 2014), we did not find evidence for reconsolidation triggered by the reactivation session restricted to ethanol context in ethanol-free state since CHX post-reactivation fails to reduce conditioned preference. Furthermore, injecting a low dose of ethanol before reactivation session restricted to ethanol-paired context was also ineffective to induce reconsolidation. However, in other studies using similar reactivation conditions but the same drug dose as used for conditioning, reconsolidation was triggered with both morphine- and cocaine-induced CPP (Valjent et al., 2006; Taubenfeld et al., 2010; Robinson & Franklin, 2007; but see Yim et al., 2006).

Re-exposure to CS, in absence of US, is a frequently used strategy to induce memory reconsolidation, a condition shared with extinction learning protocols (Vaverková et al., 2020). The main difference between these processes is the extent of re-exposure, which is based on trace dominance theory (Eisenberg et al., 2003; Suzuki et al., 2004). Longer or repeated re-exposures to CS lead to extinction learning (Cassini et al., 2017; Li et al., 2015; Suzuki et al., 2004), creating a new memory trace where CS (contextual cues) no longer predict the outcome (drug; Bouton, 2004). Conversely, short re-exposures to CS could trigger reconsolidation, allowing an original memory to be updated (Cassini et al., 2017; Goltseker et al., 2020 Suzuki et al., 2004). Whereas disruption of extinction results in high conditioned response at a subsequent test session, impairment of reconsolidation leads to its reduction (Cassini et al., 2017; Suzuki et al., 2004). Moreover, there is a limbo in between these two-memory processes, in which neither extinction nor reconsolidation takes place, and memory is only retrieved (Cassini et al., 2017; Flavell et al., 2013; Merlo et al., 2018).

In our results, the fact that re-exposure to the entire apparatus or only to ethanol-paired context in Experiments 2-5 reduced the conditioned preference in control groups (injected with saline) could argue in favor that extinction learning was in progress. However, it is unlikely that the dominant trace in the reactivation sessions was extinction, for two reasons. First, it is well established that extinction learning depends upon synthesis of novel proteins (Cassini et al., 2017; Myers & Davis, 2002; Suzuki et al., 2004; but see Lattal & Abel, 2001) and, in our experiments, CHX did not prevent the decay of conditioned preference. Furthermore, extinction learning in CPP usually requires multiple trials in separate days to reliably reduce the preference for the ethanol compartment (Bhutada et al., 2012; Pastor et al., 2011; Pastor et al., 2011; Pildervasser et al., 2014). Second, considering that longer re-exposures to CS usually induce extinction and shorter ones induce reconsolidation, it is not likely that 5 min and 3 min reactivation sessions with free access to the apparatus generate extinction, if 10 min did not. Thus, a likely hypothesis to explain our results, is that reactivation duration with free access to the apparatus (10 min, 5 min, 3min) and restricted to ethanol-context (10 min or 5 min) could be in the limbo, between inducing extinction and reconsolidation, only allowing the ethanol-related memory to be retrieved.

In line with these findings, using contextual fear conditioning, Cassini et al. (2017) observed that MK-801, an NMDA antagonist which disrupts both reconsolidation and extinction, after intermediate duration of reactivation sessions was ineffective to disrupt reconsolidation or extinction. Moreover, both groups (MK-801 and vehicle) reduced conditioned fear behavior between reactivation and test sessions, independently of reactivation duration and which memory process was engaged (reconsolidation, retrieval, or extinction).

To disrupt memory underlying conditioned responses, the amnestic treatment should be effective to impair reconsolidation (Lee et al., 2017). Many studies showed that inhibition of protein synthesis can disrupt reconsolidation in CPP (Escosteguy-Neto et al., 2016; Milekic et al., 2006; Robinson & Franklin, 2007; (Taubenfeld et al., 2010) Valjent et al., 2006; (Yu et al., 2013) and the same dose of CHX can impair reconsolidation in both CPP and fear conditioning (Milekic et al., 2006; Haubrich et al., 2015). Finally, as a positive control, we attested that the dose of CHX (100 mg/kg) was able to impair consolidation in inhibitory avoidance task (Exp. 8; Fig. 5B), and this dose also blocked reconsolidation in auditory fear conditioning in mice (Murav’eva & Anokhin, 2007). These findings support that it is unlikely that the lack of effect of CHX in our study was due to the dose used. In two CPP studies, reconsolidation was impaired only if protein synthesis was inhibited for a longer time, with two injections of treatment (immediately and 5 h after reactivation session; Milekic et al., 2006; Taubenfeld et al., 2010). This strategy was not tested in the present study because most studies showed amnestic effect with a single injection. However, we cannot rule out this possibility to disrupt reconsolidation in our protocols.

Finally, we used a deconditioning-update approach to reduce the reward motivational value of the context paired with ethanol. We used two 10 min re-exposures to ethanol-paired context with a low dose of ethanol (0.5 g/kg) aiming to induce a negative prediction error, facilitate conditioned preference memory destabilization, and update the CS-US association, reducing the reward value of context. Popik et al. (2020) conducted a study using auditory fear conditioning with four reactivation sessions, in which each tone (CS) was paired with a weaker footshock than that used in the training. This protocol was able to reduce conditioned fear through reconsolidation mechanism, and authors suggested that the memory was updated with a reduced emotional valence. However, this approach did not reduce conditioned preference for ethanol compartment in our protocol.

Ethanol consistently induced CPP in many studies (Cunningham & Prather, 1992; Macedo et al., 2018; Pildervasser et al., 2014). Curiously, even ethanol being the drug which has more negative impacts in different countries (WHO, 2016), reconsolidation research in CPP rarely focused on ethanol. To the best of our knowledge, there are only three studies with reconsolidation in ethanol-CPP in addition to ours (Font & Cunningham, 2012; Goltseker et al., 2020; Lin et al. 2014). Despite the scarcity of studies with in CPP induced by ethanol, reconsolidation with this drug has been reported using self-administration paradigms in rodents (Barak et al., 2013; Chesworth & Corbit, 2018; Millan et al., 2013; Milton et al., 2012; Puaud et al., 2018; Schramm et al., 2016; Wouda et al., 2010; Vengeliene et al., 2015; von der Goltz et al., 2009) and even in humans (Das et al., 2015, 2018a, 2018b, and 2019; Gale et al., 2020; Hon et al., 2016). This evidence suggests that it is possible to modify a well learned reward-associated memory with ethanol.

The possibility to disrupt consolidated memories or to change its motivational/emotional valence, reducing underlying behaviors, is the main therapeutic potential for reconsolidation in mental disorders, including addiction. However, the big challenge in this field is to understand how to make memory vulnerable to post-retrieval amnestic effects (Chen et al., 2021; Exton-McGuinness & Milton, 2018). Negative results in literature, including the reports in the present study, and the variability in effective reactivation sessions in CPP studies (condition and duration) reflect the complexity to induce destabilization of appetitive memories. The occurrence of destabilization-reconsolidation will depend on 1) the boundary conditions regarding features of memory itself, 2) conditions under which it is retrieved (reactivation session), 3) the dominant memory trace during retrieval, 4) the memory type and behavioral paradigm used, and 5) the amnestic treatment (dose and molecular targets). How each one of these factors work independently or synergistically should be more investigated in basic science in order to optimize the parameters during reactivation sessions that effectively trigger reconsolidation.

## Acknowledgments

We thank funding agencies (CAPES, CNPq, and AFIP) and Letícia Pichinin who helped in some experiments.

